# GluA4 enables associative memory formation by facilitating cerebellar expansion coding

**DOI:** 10.1101/2020.12.04.412023

**Authors:** Katarzyna Kita, Catarina Albergaria, Ana S. Machado, Megan R. Carey, Martin Müller, Igor Delvendahl

## Abstract

AMPA receptors (AMPARs) mediate excitatory neurotransmission in the CNS and their subunit composition determines synaptic efficacy. Whereas AMPAR subunits GluA1–GluA3 have been linked to particular forms of synaptic plasticity and learning, the functional role of GluA4 remains elusive. Here we used electrophysiological, computational and behavioral approaches to demonstrate a crucial function of GluA4 for synaptic excitation and associative memory formation in the cerebellum. Notably, GluA4-knockout mice had ∼80% reduced mossy fiber to granule cell synaptic transmission. The fidelity of granule cell spike output was markedly decreased despite attenuated tonic inhibition and increased NMDA receptor-mediated transmission. Computational modeling revealed that GluA4 facilitates pattern separation that is important for associative learning. On a behavioral level, while locomotor coordination was generally spared, GluA4-knockout mice failed to form associative memories during delay eyeblink conditioning. These results demonstrate an essential role for GluA4-containing AMPARs in cerebellar information processing and associative learning.

## Introduction

AMPA receptors (AMPARs) are essential for excitatory neurotransmission in the central nervous system (CNS). AMPARs are tetramers assembled from a combination of four subunits, GluA1–GluA4, that have distinct properties (Traynelis et al., 2010). AMPAR subunit expression shows strong regional variations (Geiger et al., 1995; Schwenk et al., 2014; Sjöstedt et al., 2020), suggesting specific functions in different types of neurons. Previous studies have revealed roles for AMPAR subunits GluA1–GluA3 in diverse forms of synaptic plasticity across the CNS (Citri et al., 2010; Gutierrez-Castellanos et al., 2017; Renner et al., 2017; Roth et al., 2020; Shi et al., 2001; Silva et al., 2019; Steinberg et al., 2006; Zamanillo, 1999), but much less is known about the function of the GluA4 subunit.

GluA4 confers rapid kinetics and large conductance to synaptic receptors (Mosbacher et al., 1994; Swanson et al., 1997). Yet, its expression is confined to few types of neurons in the adult brain (Keinänen et al., 1990; Monyer et al., 1991). Deletion of GluA4 impairs excitatory input to certain neurons in the auditory brainstem (Yang et al., 2011), the thalamus (Paz et al., 2011; Seol and Kuner, 2015), and the hippocampus (Fuchs et al., 2007). In the cerebellum, granule cells (GCs)—the most abundant neurons in the mammalian brain—heavily express GluA4 (Cathala et al., 2005; Hollmann and Heinemann, 1994; Mosbacher et al., 1994; Schwenk et al., 2014), but the significance of GluA4 for GCs and for cerebellar circuit function remains unclear.

The cerebellum plays an important role for sensorimotor integration, motor coordination, motor learning and timing, as well as cognition (Diedrichsen et al., 2019; Raymond and Medina, 2018). The underlying computations are very fast, allowing millisecond precision in the calibration and timing of movement (Heck et al., 2013; Osborne et al., 2007). This astonishing speed is achieved by the cerebellar cortex with a highly conserved network structure: Sensory and motor information enter the cerebellar cortex via mossy fibers (MFs) that contact a large number of GCs. These small, numerous neurons greatly outnumber MFs, thus providing a large expansion of coding space (Albus, 1971; Marr, 1969). GCs relay information via their parallel fibers (PF) to Purkinje cells (PCs), the sole output neurons of the cerebellar cortex. Accumulating evidence suggests that the MF→GC synapse has evolved mechanisms allowing for transmission with exceptionally high rates and precision (Delvendahl and Hallermann, 2016; DiGregorio et al., 2002; Rancz et al., 2007). However, the molecular mechanisms underlying rapid signal integration in cerebellar GCs remain enigmatic.

Here we used electrophysiological recordings, computational modeling, and behavioral analyses to study cerebellar function in adult GluA4-knockout (GluA4-KO) mice. Deletion of GluA4 resulted in a selective impairment of MF→GC transmission. Despite compensatory changes in inhibition and NMDA receptor-mediated synaptic input, the pronounced decrease of AMPAR-mediated excitation caused a severe deficit in synaptic integration during high-frequency transmission. These synaptic changes impaired pattern separation and learning performance of a feedforward network model. On a behavioral level, GluA4-KO mice displayed normal locomotor coordination, but a complete absence of eyeblink conditioning. Our findings demonstrate the importance of the GluA4 AMPAR subunit for synaptic function and associative memory formation in the cerebellum.

## Results

### Selective Impairment of Cerebellar MF→GC Synapses in GluA4-KO Mice

GluA4 shows a regional expression pattern in the CNS with strongest detection in the cerebellum (Sjöstedt et al., 2020). Within the cerebellar cortex, this AMPAR subunit has been described in Bergmann glia (Saab et al., 2012) and in GCs (Mosbacher et al., 1994), where GluA4 is the major AMPAR subunit at MF→GC synapses (Delvendahl et al., 2019). To investigate the role of GluA4 for cerebellar function, we performed whole-cell patch-clamp recordings at excitatory synapses in slices of adult WT and GluA4-KO mice (Figure 1A). Recordings were made in the anterior vermis that mainly receives sensory MF input (lobules III– IV; Giovannucci et al., 2017; Witter and De Zeeuw, 2015). GluA4-KO GCs displayed strongly diminished synaptic responses upon MF stimulation, with a ∼80% reduction in excitatory postsynaptic currents (EPSCs; 76.2 ± 7.5 pA vs. 13.4 ± 1.1 pA; p < 0.0001; Figure 1B–C; (Delvendahl et al., 2019)). Consistent with the fast kinetics of GluA4-containing AMPARs (Mosbacher et al., 1994), EPSC decay kinetics were slower in GluA4-KO GCs (1.8 ± 0.2 ms vs. 3.1 ± 0.3 ms; p < 0.001; Figure 1C). To probe the contribution of GluA4 to excitatory synapse function along the MF pathway, we investigated MF synapses onto Golgi cells (GoCs), which provide feedforward and feedback inhibition to GCs (Duguid et al., 2015). At MF→GoC connections (Figure 1D), EPSC amplitudes and decay kinetics were comparable between WT and GluA4-KO (Figure 1E–F), suggesting that GluA4 does not contribute to MF→GoC transmission.

**Figure 1.**
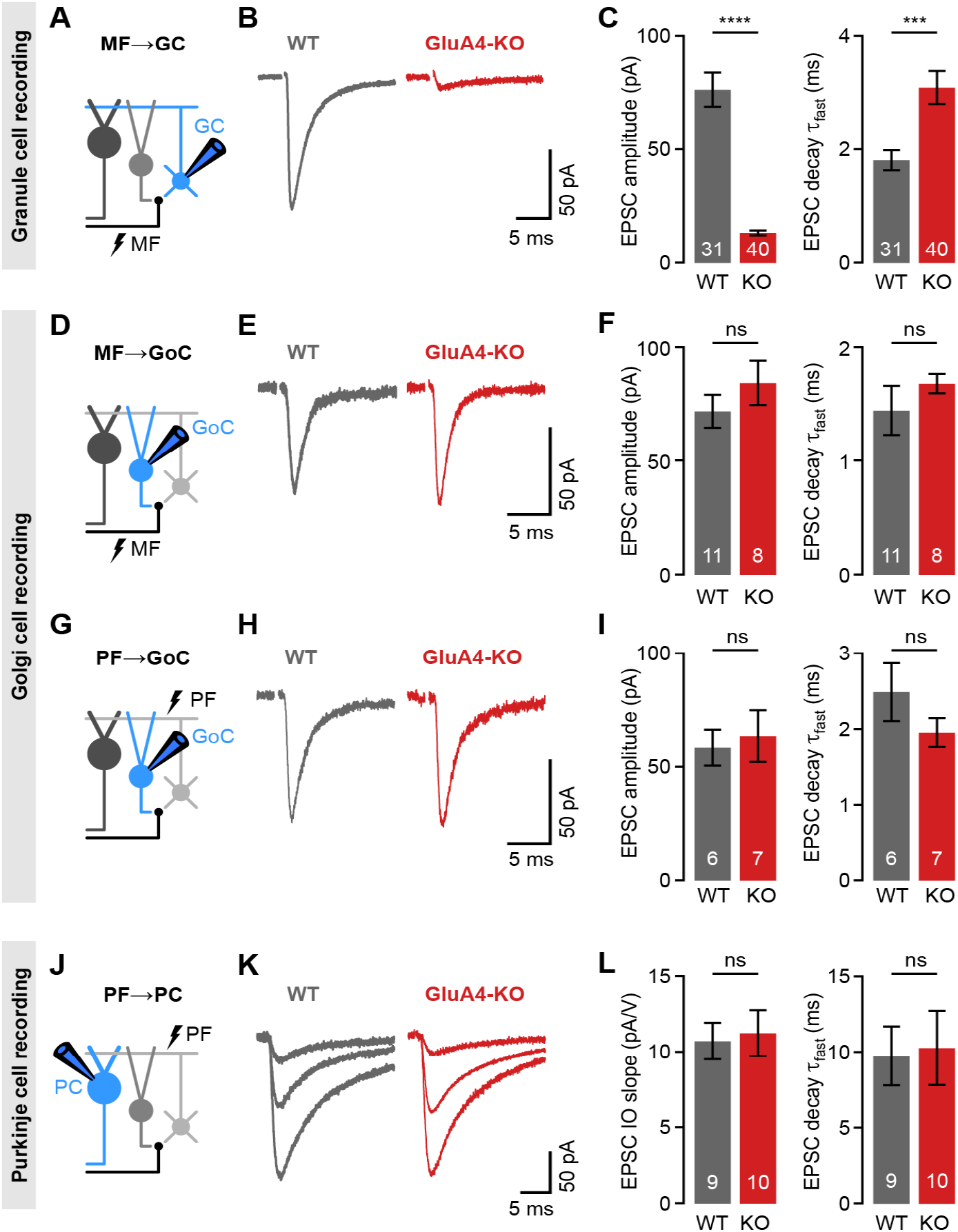
Selective Impairment of Cerebellar MF→GC Synapses in GluA4-KO Mice. **A**. Recordings from MF→GC connections. **B**. Example EPSCs recorded from WT and KO GCs by single MF stimulation. **C**. Average EPSC amplitude (left) and average EPSC fast decay time constant (right) for WT and KO MF→GC EPSCs. Data are redrawn from (Delvendahl et al., 2019). **D**. Recordings from MF→GoC connections. **E**. Example EPSCs upon MF stimulation recorded from WT and KO GoCs. **F**. Average EPSC amplitude (left) and average EPSC fast decay time constant (right) for WT and KO MF→GoC EPSCs. **G**. Recordings from PF→GoC connections. **H**. Example EPSCs recorded from WT and KO GoCs upon PF stimulation. **I**. Average EPSC amplitude (left) and average EPSC fast decay time constant (right) for WT and KO PF→GoC EPSCs. **J**. Recordings from PF→PC connections. **K**. Example EPSCs recorded from WT and KO PCs by stimulating afferent PFs with increasing stimulation strength (displayed stimulation intensities: 7, 12 and 17 V). **L**. Average linear slope of the input-output relationship (left) and average EPSC fast decay time constant (right) for WT and KO PF→PC. Data are means ± SEM.

To probe if deletion of GluA4 affects the function of GC output synapses, we first investigated PF inputs to GoCs (PF→GoC synapses; Figure 1G), which were similar between WT and GluA4-KO mice (Figure 1H–I). Consistent with the similar EPSC amplitudes, spontaneous EPSCs and paired-pulse ratios were also not altered in GoCs of GluA4-KO mice (Figure S1A–D). We next studied PF→PC synapses (Figure 1J). Stimulation in the molecular layer with increasing intensity revealed similar PF→PC EPSCs in WT and GluA4-KO (Figure 1K). The slope of the input-output relationship was comparable between genotypes (Figure 1L; Figure S1G), which indicates normal recruitment of PFs. Likewise, EPSC decay kinetics (Figure 1L), as well as spontaneous EPSC amplitudes and paired-pulse ratios were similar in WT and KO PCs (Figure S1E–H), suggesting that GluA4 does not contribute directly to PF→PC synaptic transmission. Together, the recordings from PF→GoC and PF→PC synapses indicate that PF output is functionally intact in GluA4-KO mice. Our findings demonstrate that GluA4 is indispensable for fast neurotransmission at MF→GC synapses and suggest a GC-specific impairment of excitatory transmission at the cerebellar input layer in GluA4-KO mice.

### Reduced Tonic Inhibition Enhances GC Intrinsic Excitability

Loss of GluA4 severely compromises synaptic excitation of cerebellar GCs. Changes in both excitatory and inhibitory synaptic input may influence a neuron’s intrinsic excitability (Aizenman and Linden, 2000; Desai et al., 1999; Karmarkar and Buonomano, 2006). To investigate if GluA4-KO alters the intrinsic excitability of GCs, we quantified GC spiking upon increasing tonic current injections (Figure 2A). Under control conditions, GluA4-KO GCs showed higher firing frequencies than WT, reflecting increased excitability (Figure 2B–C). Because GoC-mediated tonic inhibition influences the intrinsic excitability of GCs both in slices (Brickley et al., 2001; Hamann et al., 2002; Mitchell and Silver, 2003; Rothman et al., 2009) and *in vivo* (Chadderton et al., 2004), we next probed GC firing in the presence of the GABA_A_ receptor (GABA_A_R) blocker bicuculline. Interestingly, bicuculline increased spiking only in WT GCs, to a level that was comparable to GluA4-KO GCs (ANOVA: current × genotype × drug: F = 5.34, p = 0.02; Figure 2B–C). Blocking inhibition increased the gain and reduced rheobase in WT (Figure 2D), consistent with enhanced intrinsic excitability. By contrast, bicuculline had no effect on gain and rheobase current in GluA4-KO GCs (ANOVA gain: genotype × drug: F = 4.24, p = 0.041; rheobase: genotype × drug: F = 2.00, p = 0.16; Figure 2D), indicating that tonic inhibition is reduced in these mice.

**Figure 2.**
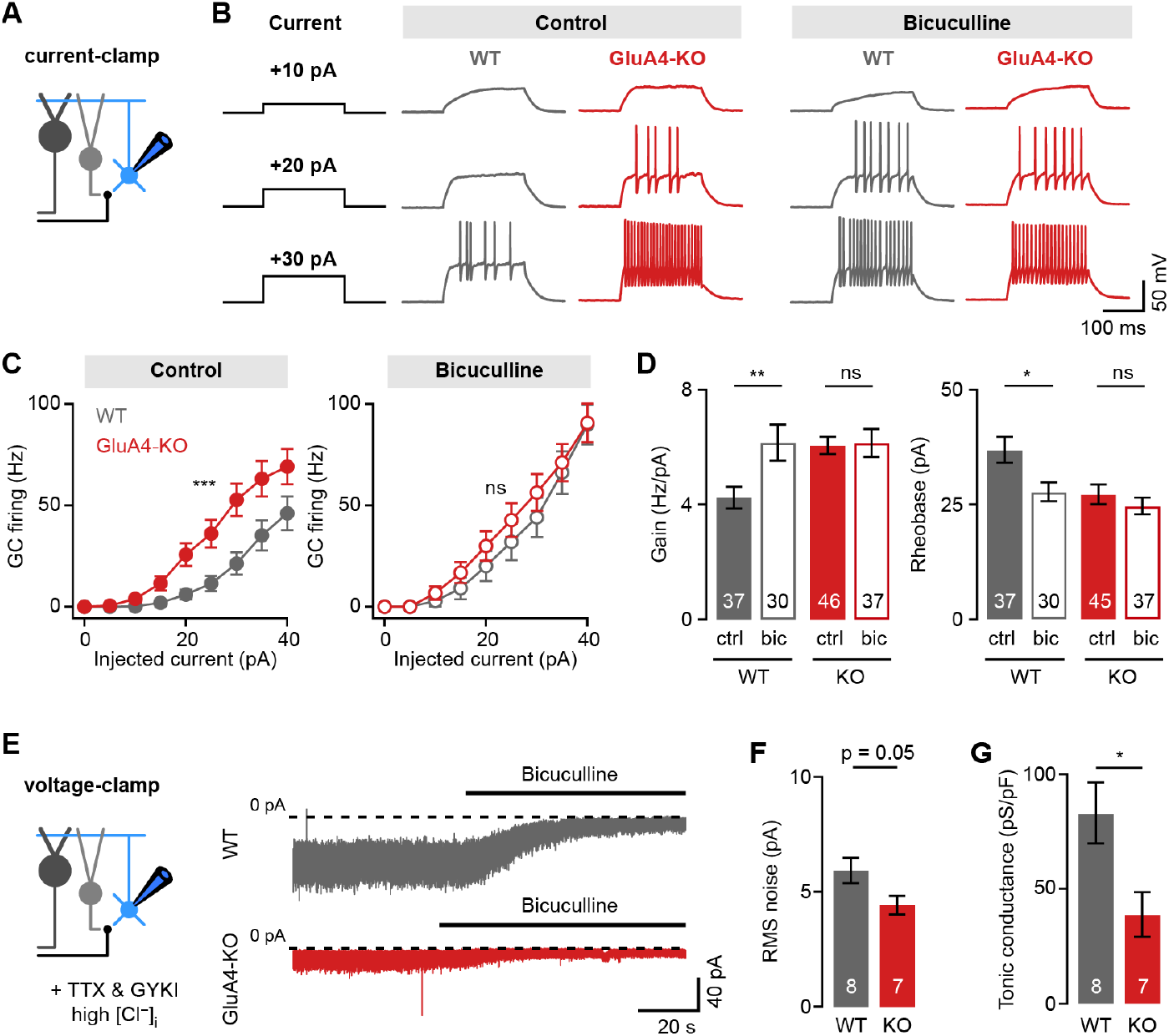
Reduced Tonic Inhibition Enhances GC Intrinsic Excitability. **A**. Current-clamp recordings from GCs. **B**. Example responses to somatic current injection of indicated amplitude for control and in the presence of 10 µM bicuculline. **C**. Average GC firing frequency plotted vs. injected current for WT and KO in control and bicuculline. **D**. Average gain and rheobase, calculated from the frequency-current curves in (**C**). In the presence of bicuculline, KO GCs had gain and rheobase values comparable to WT. **E**. Recording of tonic holding current. GABAergic currents were isolated using 20 µM GYKI-53655 and high intracellular Cl^−^. Right: Example recordings of bicuculline wash-in (10 µM) for WT and KO GCs. **F**. Left: Average root-mean-square noise before wash-in of bicuculline (left) and average tonic conductance density (right) for WT and GluA4-KO. Tonic conductance was calculated from the bicuculline-sensitive current. Data are means ± SEM.

Persistent activation of *δ*- and *α*6-containing extrasynaptic GABA_A_Rs causes a tonic inhibitory conductance in cerebellar GCs (Brickley et al., 2001; Hamann et al., 2002; Stell et al., 2003) that also influences their excitability (Figure S2A–C; (Mitchell and Silver, 2003; Rudolph et al., 2020)). We directly investigated tonic inhibition by isolating inhibitory conductance in WT and GluA4-KO GCs and bath-applying bicuculline. Indeed, GluA4-KO GCs had a smaller root-mean-square noise of the baseline holding current and smaller bicuculline-sensitive conductance compared with WT (WT: 83.0 ± 13.3 pS/pF, KO: 39.0 ± 9.6 pS/pF, p = 0.022; Figure 2E–F). These findings demonstrate that tonic GABA_A_R-mediated inhibition is reduced in GluA4-KO GCs, in line with the marginal effect of bicuculline on intrinsic excitability of GluA4-KO GCs. By contrast, the amplitudes and frequencies of spontaneous inhibitory postsynaptic currents (sIPSCs) were not different between WT and GluA4-KO GCs (Figure S2E), indicating that phasic inhibition is not altered. To verify the specificity of our findings, we also studied excitability in GCs of heterozygous GluA4 mice (GluA4-HET), which have reduced GluA4 levels (Figure S5A) but no change in MF→GC EPSC amplitudes (Figure S5E–F; (Delvendahl et al., 2019)). We did not observe indications of altered inhibition in GluA4-HET GCs (Figure S5B–D). Thus, impaired excitatory synaptic input to GCs is accompanied by a modulation of tonic inhibition, resulting in increased intrinsic excitability.

### Shunting Inhibition Controls EPSP Size in Cerebellar GCs

Tonic GABA_A_R activation leads to shunting inhibition of GCs that may affect their responses to excitatory input (Brickley et al., 2001; Mitchell and Silver, 2003). The reduction of tonic inhibition in GluA4-KO GCs is therefore expected to impact excitatory postsynaptic potentials (EPSPs). To assess the impact of shunting inhibition in WT and GluA4-KO GCs, we recorded MF→GC EPSCs and EPSPs in blocker-free solution (‘control’, Figure 3A). As expected, EPSC amplitudes were strongly diminished in GluA4-KO GCs, as were EPSP amplitudes (Figure 3B–C). Interestingly, the amplitude reduction relative to WT was smaller for EPSPs than for EPSCs (69.4 ± 2.1% vs. 80.6 ± 1%, p < 0.001; Figure 3C). To test if reduced tonic inhibition underlies this effect, we recorded EPSPs in the presence of bicuculline, which increased EPSP amplitudes in WT, but not in GluA4-KO GCs (ANOVA drug × genotype: F = 8.00, p = 0.005; Figure 3D–E). We also observed that bicuculline enhanced EPSP amplitudes during high-frequency train stimulation in WT, but not in GluA4-KO GCs (ANOVA drug × genotype: F = 46.88, p < 0.001; Figure 3F–G). In contrast, bicuculline had no differential effect on EPSC amplitudes (ANOVA drug × genotype: F = 0.35, p = 0.56; Figure S2F–G). These results demonstrate that shunting inhibition reduces postsynaptic depolarization in response to excitatory input in WT, but not in GluA4-KO GCs. We conclude that the reduced tonic inhibition upon loss of the major AMPAR subunit at MF→GC synapses augments synaptic depolarization in GluA4-KO GCs.

**Figure 3.**
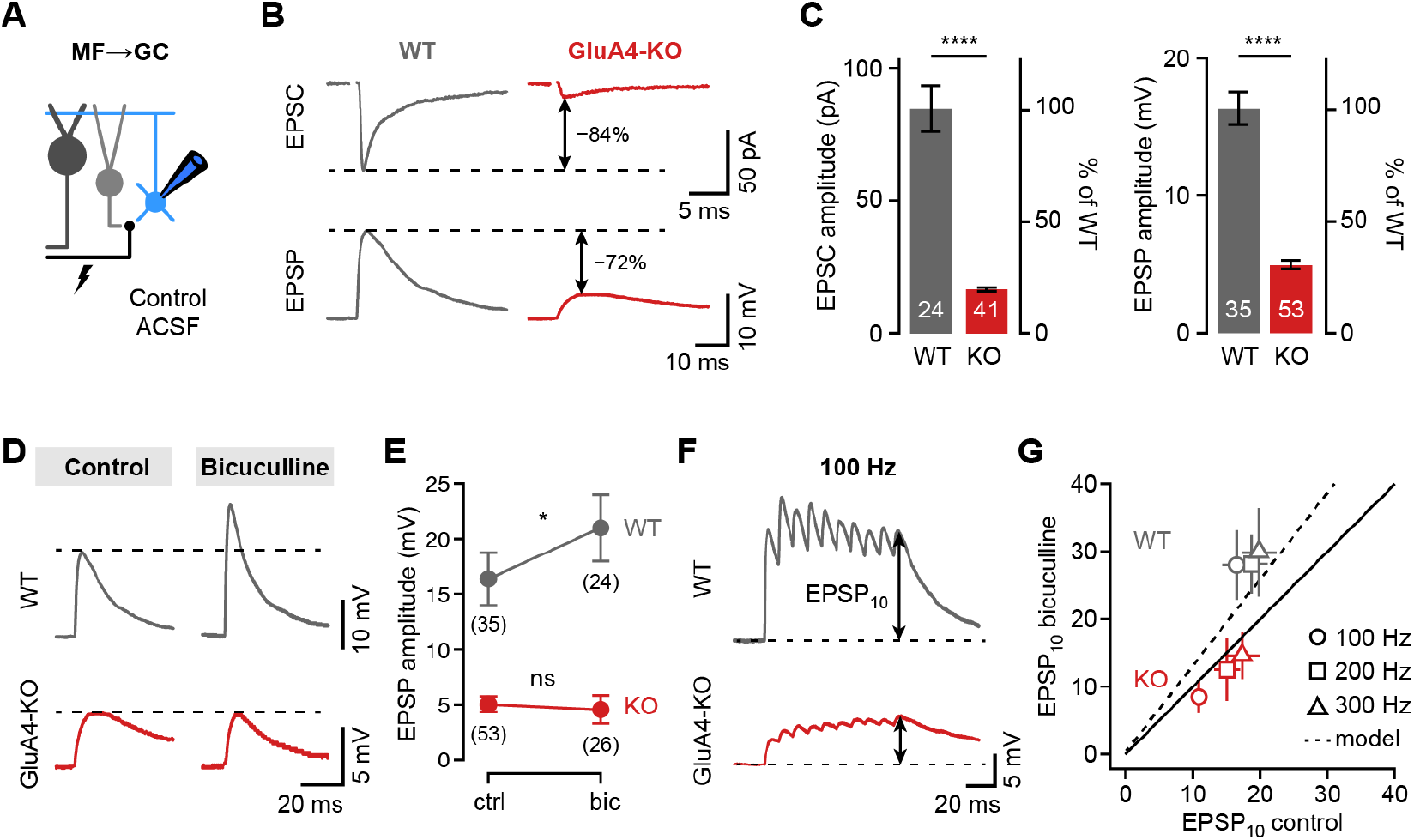
Shunting Inhibition Controls EPSP Size in Cerebellar GCs. **A**. Recordings from MF→GC connections in voltage- or current-clamp with blocker-free ACSF (‘control’). **B**. Example EPSC and EPSP recordings from the same cell in WT and KO. Arrows indicate reduction compared to WT. **C**. Average EPSC and EPSP amplitudes for WT and KO. Note the difference in relative reduction of EPSPs compared to EPSCs. **D**. Example EPSP recordings for control and in the presence of 10 µM bicuculline. **E**. Average EPSP amplitude for WT and KO. Error bars represent 95% CI. **F**. Example MF→GC EPSP trains (10 stimuli, 100 Hz). **G**. Average EPSP_10_ in the presence of bicuculline vs. control for the indicated stimulation frequencies. Black line represents unity and dashed line the prediction of an integrate-and-fire model with or without tonic inhibition (cf. Figure S2). Data in (**C**) and (**G**) are means ± SEM.

### Deletion of GluA4 Impairs GC Spike Fidelity and Precision during High-Frequency Transmission

How do the changes in excitatory input and intrinsic excitability influence GC firing in response to MF input? To address this question, we performed current-clamp recordings from GCs in combination with high-frequency MF→GC stimulation (Figure 4A) at a membrane potential of −70 mV, corresponding to the average resting membrane potential of GCs *in vivo* (Chadderton et al., 2004; Powell et al., 2015). WT GCs showed increasing firing frequencies upon 100–300 Hz stimulation (Figure 4B–C). We also observed MF-stimulation-evoked spikes in GluA4-KO GCs, albeit at reduced frequency (ANOVA genotype: F = 23.48; p < 0.0001; Figure 4B) and in fewer cells (Figure 4C). The GC spikes elicited by high-frequency MF→GC stimulation were not only reduced in number, but also occurred with a longer delay compared to WT (ANOVA genotype: F = 26.26; p < 0.0001; Figure 4D–E). Hence, GluA4-KO GCs need to summate more EPSPs to reach spike threshold (Figure S3D), leading to reduced reliability and impaired temporal precision of GC spikes. Spike output was not altered in GCs of GluA4-HET mice (Figure S5H–I). Together, these data show a strong reduction of spiking frequency and temporal precision in GluA4-KO GCs upon high-frequency synaptic input, and establish that GluA4-containing AMPARs control the timing and reliability of EPSP-spike coupling at the cerebellar input layer.

**Figure 4.**
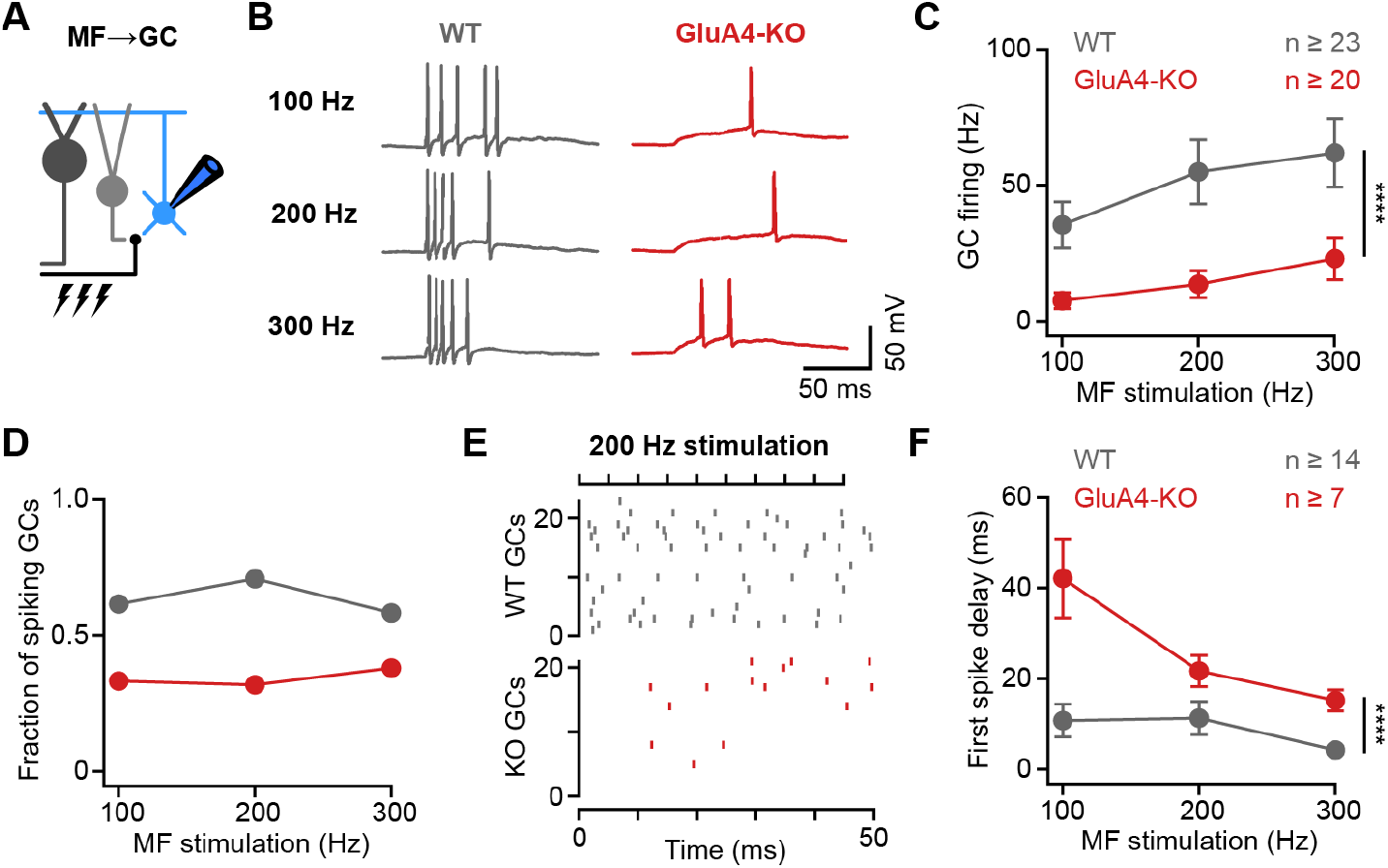
Deletion of GluA4 Impairs GC Spike Fidelity and Precision during High-Frequency Transmission. **A**. Recordings from MF→GC connections in current-clamp. **B**. Example voltage recordings from a WT and GluA4-KO GC upon MF stimulation with 10 stimuli at increasing frequencies (indicated). **C**. Average GC firing frequency upon MF stimulation with 100–300 Hz frequency. GCs were held at −70 mV. **D**. Fraction of GCs showing spikes upon MF stimulation with 100–300 Hz frequency. **E**. Spike raster plots for 200 Hz MF stimulation in WT and GluA4-KO. **F**. Average first spike delay calculated from stimulation onset plotted versus MF stimulation frequency. Data are means ± SEM.

### NMDARs and Reduced Inhibition Support Spiking in GluA4-KO GCs

Given the strong reduction of MF→GC EPSCs (Figure 1), the occurrence of spikes upon repetitive stimulation in more than 30% of GluA4-KO GCs is surprising. We previously provided evidence for enhanced presynaptic glutamate release from GluA4-KO MF boutons (Delvendahl et al., 2019), which might increase the non-AMPAR component of MF→GC transmission. Indeed, isolated NMDA receptor (NMDAR)-mediated MF→GC EPSCs were larger in GluA4-KO than in WT GCs (27.5 ± 2.1 pA vs. 37.2 ± 2.1 pA; p = 0.0029; Figure 5A–B). Because NMDAR activation can significantly contribute to GC excitation during repetitive input (D’Angelo et al., 1995), we recorded isolated AMPAR- and NMDAR-EPSCs upon high-frequency train stimulation (100 Hz, 20 stimuli). As expected, AMPAR-EPSCs were strongly decreased in GluA4-KO GCs (Figure 5C), leading to a pronounced reduction of cumulative EPSC charge during the train (3.4 ± 0.2 pC vs. 0.9 ± 0.1 pC; p < 0.001; Figure 5D). By contrast, NMDAR-EPSCs and cumulative EPSC charge were increased in GluA4-KO GCs (5.6 ± 0.4 pC vs. 8.7 ± 0.9 pC; p = 0.013; Figure 5D).

**Figure 5.**
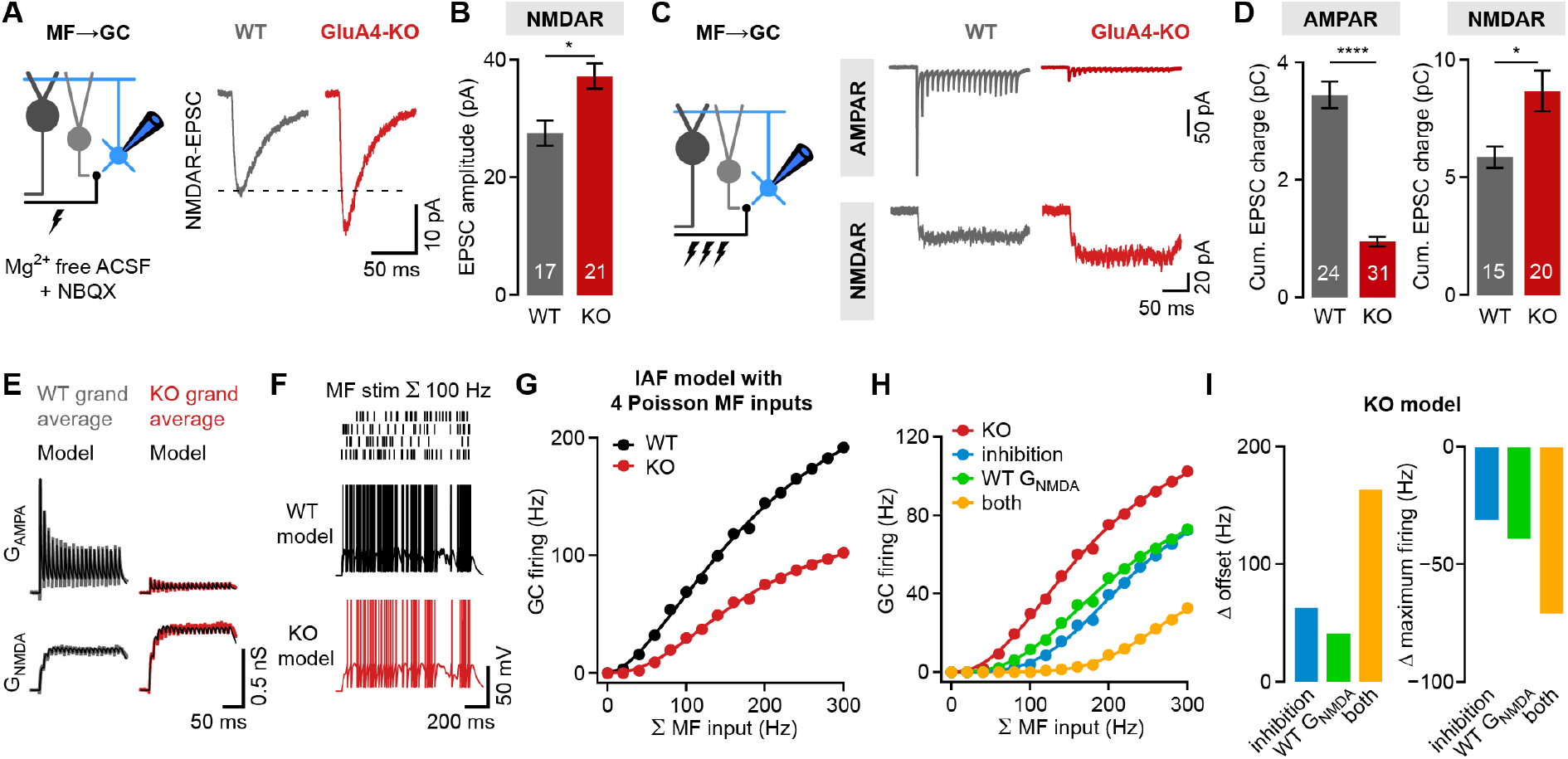
NMDARs and Reduced Inhibition Support Spiking in GluA4-KO GCs. **A**. Left: Recordings of isolated NMDAR-EPSCs at MF→GC connections. Right: Example NMDAR-mediated MF→GC EPSCs. **B**. Average NMDAR-EPSC amplitude (right) for WT and GluA4-KO GCs. **C**. Top: Example AMPAR-EPSC 100-Hz train recordings (20 stimuli) for WT and KO. Bottom: Example NMDAR-mediated 100-Hz trains. **D**. Average cumulative charge for AMPAR-mediated (left) and NMDAR-mediated EPSCs (right) for both genotypes. **E**. Grand average of AMPAR- and NMDAR-conductance trains (20 stimuli, 100 Hz) recorded from WT (left) and GluA4-KO GCs (right). Data are overlaid with synaptic conductance models for WT and GluA4-KO. **F**. Top: Four independent Poisson-distributed MF spike trains with sum of 100 Hz (top) and simulation results for WT (black) and KO (red) (bottom). **G**. Average WT and KO model firing frequency versus MF stimulation frequency. Lines are fits with a Hill equation. **H**. Results for KO model compared with simulations of KO GCs with tonic inhibition (blue), with WT G_NMDA_ (green) or both (yellow). **I**. Reduced inhibition and enhanced G_NMDA_ increase the maximum firing frequency (left) and reduce the offset (right) of the KO model GC output. Data are means ± SEM.

To investigate if reduced tonic inhibition and enlarged NMDAR-EPSCs support spiking in GluA4-KO GCs, we employed computational modeling. We fit the recorded MF→GC AMPAR- and NMDAR-conductance trains (G_AMPA_ and G_NMDA_, respectively; Figure 5E) to model synaptic input of an integrate-and-fire neuron (Rothman et al., 2009). Simulations reproduced the experimentally observed GC firing and first spike delay well for WT and GluA4-KO upon fixed frequency MF input with a binomial short-term plasticity model (Rothman and Silver, 2014) (Figure S3B– C). To probe the impact of tonic inhibitory conductance and G_NMDA_ enhancement, we simulated four independent, Poisson-distributed MF inputs (Figure 5F) covering the wide range of MF frequencies observed *in vivo* (Arenz et al., 2008; van Kan et al., 1993). The model with GluA4-KO G_AMPA_ and G_NMDA_ and reduced tonic conductance displayed lower firing frequencies over the entire range of MF inputs (Figure 5G; Figure S4A). Compared with WT, the KO model had a ∼44% reduced maximum firing frequency (186.1 Hz vs. 336.4 Hz) and a ∼17 Hz increased offset, similar to the experimental findings (Figure 4C). As in the current-clamp recordings (Figure 4F), spikes occurred with a longer delay in the KO GC model over all simulated MF frequencies (Figure S4D).

To understand the individual contributions of reduced tonic inhibition and enhanced G_NMDA_ to spiking in KO GCs, we either added an inhibitory conductance of 0.16 nS (calculated from Figure 2F) or used the WT G_NMDA_ in the KO model. In both scenarios, firing frequencies were reduced in relation to the KO model, with each mechanism contributing ∼20% to the maximum spiking frequency of the KO model (Figure 5H–I, Figure S4A–B). Combining inhibition and WT G_NMDA_ reduced the firing rate by ∼40% (Figure 5H–I), implying a synergistic, linear interaction of the two.

We next asked how the experimentally observed changes in GluA4-KO GCs influence synaptic information transfer. Analysis of pre- and postsynaptic spike trains revealed that the strongly reduced G_AMPA_ of the KO model compromises the synchrony of MF and GC spikes (∼2-fold reduction in SPIKE synchronization; Figure S4F; (Mulansky et al., 2015)), and that the reduced tonic inhibition and G_NMDA_ enhancement in GluA4-KO facilitate MF→GC synaptic information transfer at low input frequencies, albeit at a reduced level (Figure 5G–H, Figure S4E–F). Thus, although MF→GC AMPAR-EPSCs are strongly reduced, synaptic and non-synaptic mechanisms cooperatively counteract the impaired spiking output of GluA4-KO GCs. However, both mechanisms are insufficient to compensate for the pronounced impairment of synaptic excitation caused by the deletion of GluA4.

### Impaired Pattern Separation in a Feedforward Model of the Cerebellar Input Layer

The GC layer is ideally suited for sparsening and expanding of MF inputs (Albus, 1971; Marr, 1969). This transformation of neural coding is thought to enable effective pattern separation and facilitate associative learning in downstream circuits (Cayco-Gajic and Silver, 2019). To address how the reduced synaptic input in GluA4-KO GCs affects information processing in the GC layer, we employed computational modeling of a feedforward MF→GC network with constrained connectivity and biophysical properties (Figure 6A–B; (Cayco-Gajic et al., 2017)). We systematically varied the fraction of active MFs and their spatial correlation, and analyzed how the GC network transforms MF activity patterns (Cayco-Gajic et al., 2017). To assess the impact of reduced synaptic excitation, we scaled model synaptic conductances and tonic inhibition according to our GluA4-KO data (see Methods). Both control and KO models showed a similar increase in spatial sparseness of GC activity patterns compared to MF activity patterns (Figure 6C), indicating that deletion of GluA4 did not affect population sparsening. In addition to sparsening, an important characteristic of the GC layer is an expansion of coding space, which promotes pattern separation by increasing the distance between activity patterns. To investigate expansion recoding, we analyzed the size of the distribution of activity patterns by calculating the total variance of GC population activity, normalized to the total variance of the MF population activity (Cayco-Gajic et al., 2017). The normalized total variance—reflecting the expansion of coding space—was strongly reduced in the GluA4-KO model (Figure 6D), suggesting that the increased spiking threshold of GluA4-KO GCs leads to a strong reduction of coding space at the cerebellar input layer.

**Figure 6.**
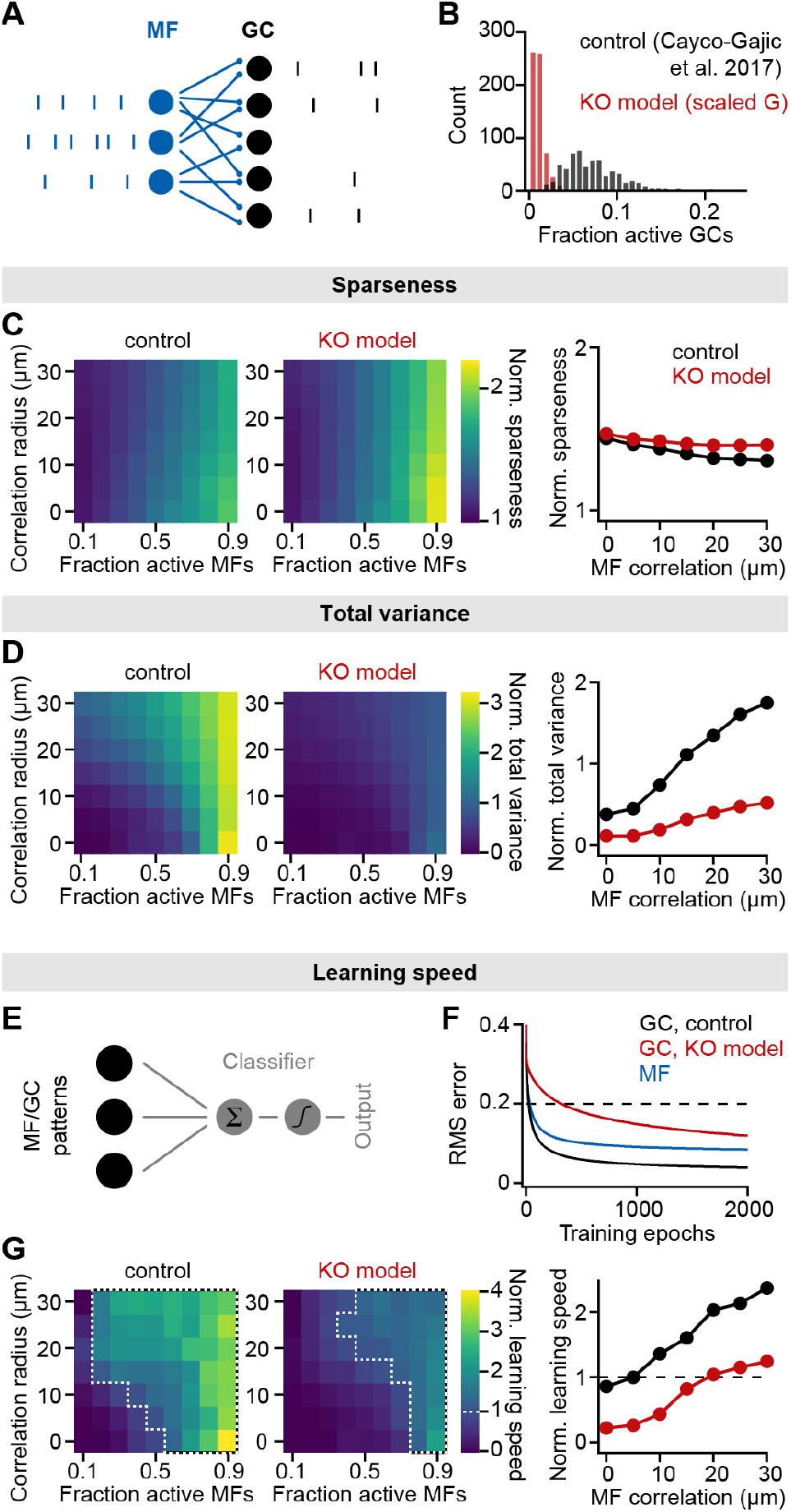
Impaired Pattern Separation in a Feedforward Model of the Cerebellar Input Layer. **A**. Schematic of the feedforward network model. The model comprises 187 MFs that are connected to 487 GCs with 4 synapses per GC. The network is presented with different MF activity patterns, which produce GC spike patterns. **B**. Histogram of active GCs. Scaling synaptic and tonic conductances according to GluA4-KO data (“KO model”, red) leads to strong reduction of active GCs compared to the model from Cayco-Gajic et al. (2017) (“control”, black). **C**. Left: Normalized population sparseness (normalized to MF sparseness) plotted for different MF correlation radii and fraction of active MFs of both models. Right: Median normalized sparseness versus MF correlation radius. **D**. Same as **C**, but for total variance. **E**. Schematic of learning. MF or GC activity patterns are used to train a perceptron decoder to classify these patterns into 10 random classes. **F**. Root-mean-square (RMS) error of the perceptron classification for MF input patterns (blue), GC input patterns (black) or GC patterns with scaled conductance (red). Dashed line indicates cutoff for learning speed quantification. **G**. Left: Normalized learning speed (normalized to learning using MF patterns) plotted for different MF correlation radii and fraction of active MFs of both models. White dashed borders indicate areas of faster learning with GC activity patterns than with MF patterns. Right: Median normalized learning speed versus MF correlation radius. Dashed line indicates faster GC learning.

To address the consequences of alterations in sparsening and expansion recoding for learning performance, we trained a perceptron to classify either MF or GC activity patterns into ten random classes (Figure 6E), and analyzed learning speed using GC patterns normalized to MF patterns (Cayco-Gajic et al., 2017) (Figure 6F). Learning speed was reduced over the full range of model parameters when using the GluA4-KO synaptic conductances (Figure 6G). Notably, GC activity accelerated learning only at high correlation levels and active fraction of MFs, in contrast to the previously published model (Cayco-Gajic et al., 2017) (white borders in Figure 6G). This result is consistent with our current-clamp experiments, where higher MF stimulation frequencies elicited GC spikes in GluA4-KO mice (Figures 4C and S3A). Moreover, the total variance (Figure 6D)—but not population sparseness—closely predicted the learning speed, suggesting that the strong decrease of MF→GC transmission impairs pattern separation by reducing the coding space. Thus, GluA4 speeds learning by improving pattern separation performance in a feedforward MF→GC network model without affecting GC population sparseness.

### Normal Locomotor Coordination of GluA4-KO Mice during Overground Walking

We next investigated the functional consequences of the extreme reduction in MF→GC transmission in GluA4-KO mice for cerebellum-dependent behaviors. Cerebellar dysfunction often leads to impairments in motor coordination, including gait ataxia during walking (Morton and Bastian, 2007). We analyzed the locomotor behavior of GluA4-KO mice using the LocoMouse system (Machado et al., 2015) (Figure 7A). GluA4-KO mice walked similarly to size-matched WT littermates without obvious gait ataxia (Supplementary Video 1). Quantitative analysis of locomotor coordination (Machado et al., 2015, 2020) (Figure 7B–C) revealed that their locomotor behavior was largely intact. GluA4-KO mice tended to walk more slowly than controls, but across walking speeds, stride lengths were comparable in mice of both genotypes (Figure 7D–E, Figure S6A–B). Paw trajectories measured by continuous forward velocity were also intact (Figure 7F–G, Figure S6E–F). Vertical paw motion was also largely normal (Figure 7H–I), although there was a small degree of hyperflexion across walking speeds (Figure S6F–G). We also explored interlimb coordination, which is often disrupted by cerebellar damage (Machado et al., 2015, 2020; Morton and Bastian, 2007), during normal walking. We did not observe any differences between WT and GluA4-KO mice, with both genotypes displaying a symmetrical trot pattern across walking speeds (Figure 7J–K). Exhaustive analysis of the movements of individual limbs, interlimb and body coordination, and comparison with other mouse models of cerebellar ataxia confirmed the grossly normal locomotor phenotype of GluA4-KO mice (Figures S6–7). Moreover, GluA4-HET mice showed normal overall locomotion performance (Figure S8A–H). Thus, locomotor coordination was spared in GluA4-KO mice despite the pronounced impairment of synaptic excitation at the cerebellar input layer.

**Figure 7.**
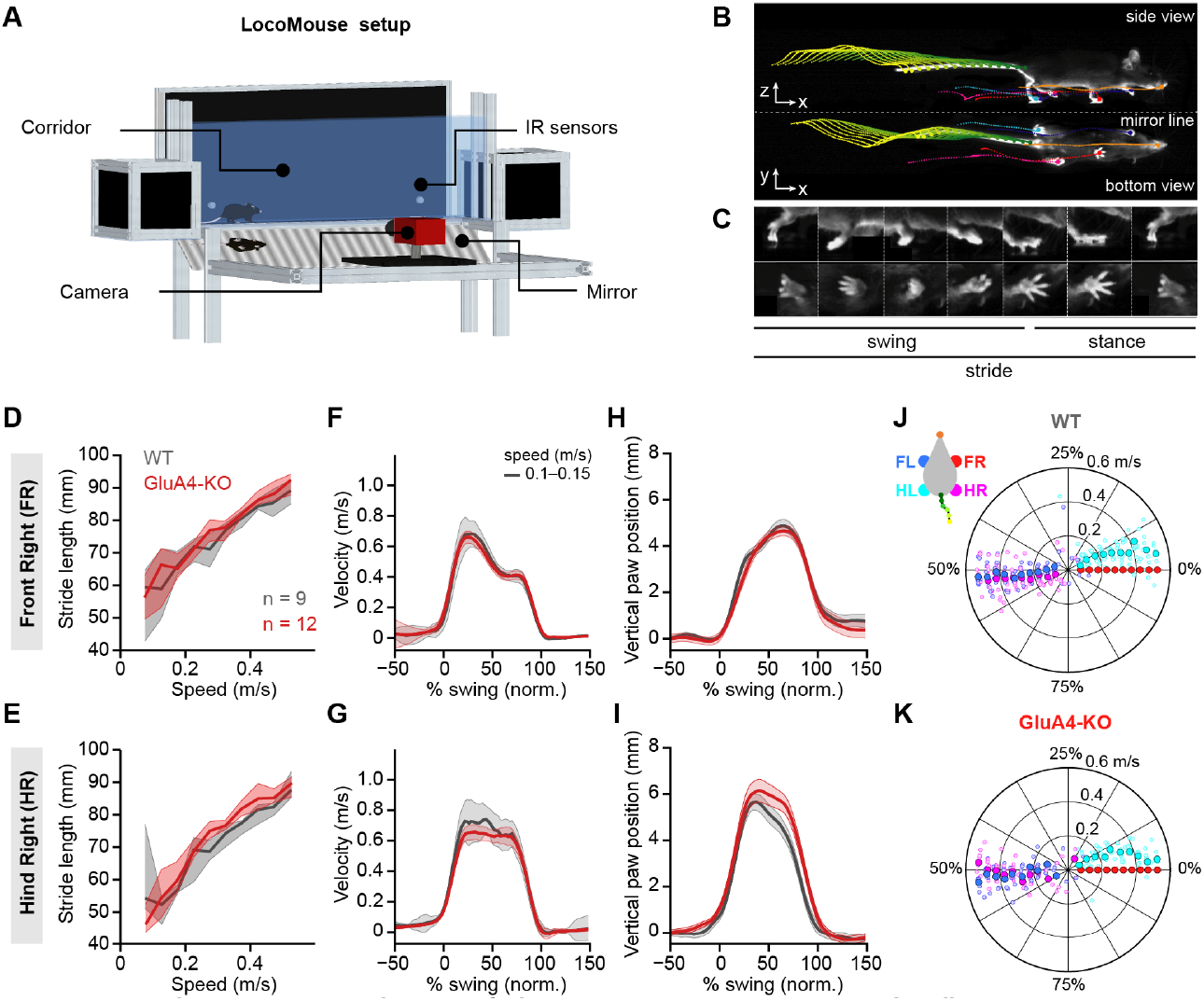
Normal Locomotor Coordination of GluA4-KO Mice During Overground Walking. **A**. LocoMouse apparatus. **B**. Continuous tracks (in x, y, z) for nose, paws and tail segments obtained from LocoMouse tracking (Machado et al., 2015) are plotted on top of the frame. **C**. Individual strides were divided into swing and stance phases for further analysis. **D–E**. Stride length of FR paw (**D**) and HR paw (**E**) vs. walking speed for GluA4-KO mice (red) and WT animals (gray). For each parameter, the thin lines with shadows represent median values ± 25th, 75th percentiles. **F–G**. Average instantaneous forward (x) velocity of FR paw (**F**) and HR paw (**G**) during swing phase. **H–I**. Average vertical (z) position of FR paw (**H**) and HR paw (**I**) relative to ground during swing. The shaded area indicates SEM across mice. **J–K**. Polar plots indicating the phase of the step cycle in which each limb enters stance, aligned to stance onset of FR paw (red circle). Distance from the origin represents walking speed. **J**. WT mice; **K**. GluA4-KO mice. Circles show average values for each animal.

### GluA4 is Required for Cerebellum-Dependent Associative Memory Formation

Influential theories of cerebellar function have proposed that pattern separation at the input layer facilitates learning within the cerebellar cortex (Albus, 1971; Cayco-Gajic and Silver, 2019; Marr, 1969). Thus, while the strong reduction of GC synaptic excitation did not impair locomotor coordination of GluA4-KO mice, it might specifically affect cerebellum-dependent learning. We therefore examined delay eyeblink conditioning in GluA4-KO mice, a cerebellum-dependent form of associative learning that involves MF inputs conveying the conditioned stimulus (CS) (Albergaria et al., 2018; De Zeeuw and Yeo, 2005; Mauk, 1997). A conditioning light pulse was repeatedly paired with an unconditioned air-puff stimulus (US) to the eye as described previously (Albergaria et al., 2018) (Figure 8A). Over training sessions, WT animals acquired a conditioned eyelid closure that preceded the US (Figure 8B). WT mice displayed a continuous increase in the percentage of trials with a conditioned response (CR; Figure 8C), demonstrating associative memory formation. Strikingly, GluA4-KO mice failed to develop CRs over the ten training sessions (Figure 8C, top), indicating a lack of associative learning in the eyeblink conditioning task. GluA4-HET animals, on the other hand, were able to acquire associative memories similarly to WT (Figure S8). Analysis of CS-only trials revealed that, in contrast to WT animals, which had developed well-timed CRs by the end of training, GluA4-KO mice failed to show any learned eyelid closures in response to the CS (Figure 8D–E, top). To rule out the possibility that impaired vision was contributing to this effect (Gründer et al., 2000; Qin and Pourcho, 1999), we repeated the eyeblink conditioning experiments in a new set of mice using whisker stimulation as a CS. Again, conditioned responses were absent in GluA4-KO mice (Figure 8B–E, bottom), demonstrating that GluA4 is required for eyeblink conditioning independent of CS modality. Thus, GluA4 appears to be dispensable for the control of coordinated locomotion (Figure 7), but is required for normal cerebellum-dependent associative learning.

**Figure 8.**
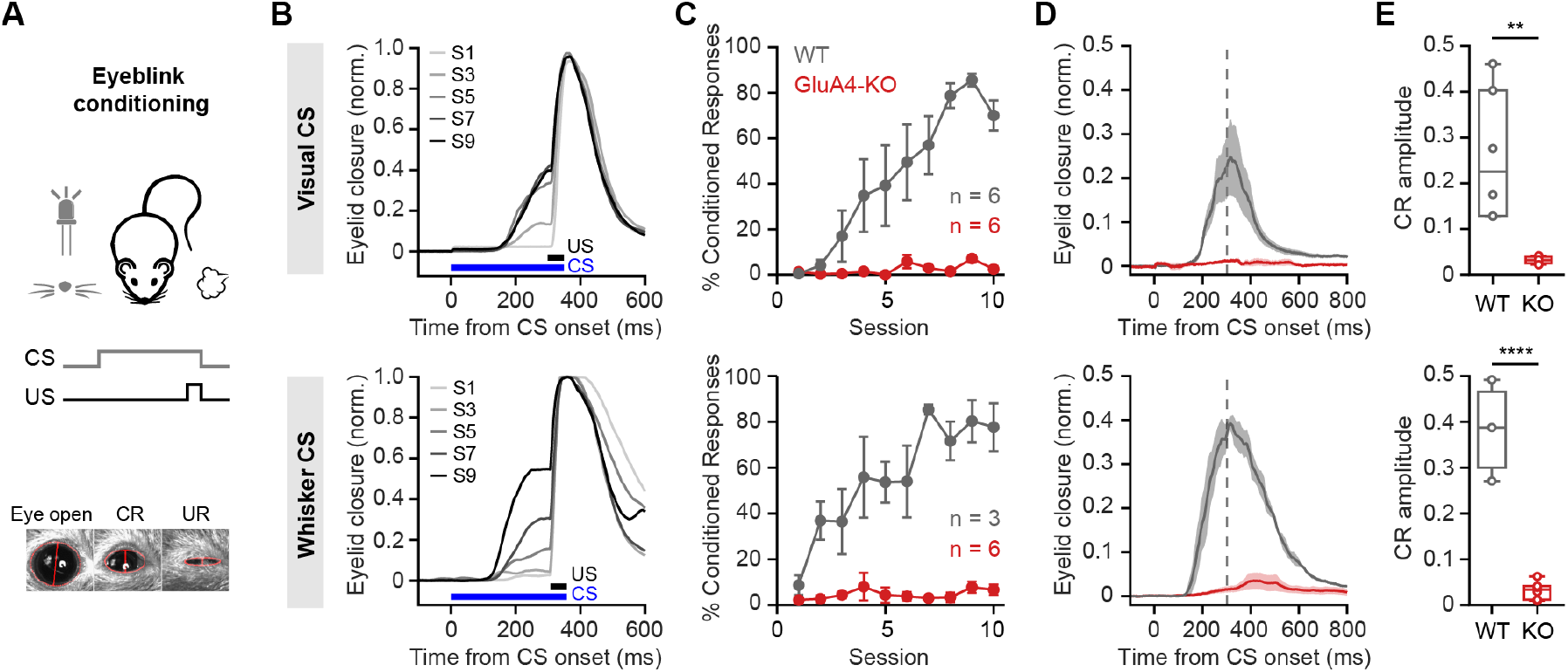
GluA4 is Required for Cerebellum-Dependent Associative Memory Formation. **A**. Top: Schematic of the eyeblink conditioning paradigm, where a CS (a white LED or a whisker vibratory stimulus) is repeatedly paired and co-terminates with an eyeblink-eliciting US (a puff of air delivered to the eye). Bottom: Example video frames (acquired at 900 fps under infrared light) illustrate automated extraction of eyelid movement amplitude. **B**. Average eyelid closure across 9 learning sessions (S1–S9) of two representative WT animals, using a visual CS (top) or a whisker CS (bottom). Each trace represents the average of 100 paired trials from a single session. **C**. Average %CR learning curves of WT (gray) and GluA4-KO (red) mice trained with either a visual CS (top) or a whisker CS (bottom). Error bars indicate SEM. **D**. Average eyelid traces of CS-only trials from the last training session of WT and GluA4-KO animals, trained with either a visual CS (top) or a whisker CS (bottom). Shadows indicate SEM. Vertical dashed line represents the time that the US would have been expected on CS+US trials. **E**. Average CR amplitudes from the last training session of WT and GluA4-KO animals, trained with either a visual CS (top) or a whisker CS (bottom). Box indicates median and 25th–75th percentiles; whiskers extend to the most extreme data points.

## Discussion

In the present study, we elucidated the functional significance of GluA4 in the cerebellum. Deletion of this AMPAR subunit strongly impaired both excitatory transmission at the cerebellum input layer and associative learning in adult mice. Despite compensatory enhancements in intrinsic excitability and NMDAR-mediated transmission that contributed to the residual function of GluA4-KO GCs, their spike output remained reduced. Computational modeling suggests that GluA4 facilitates pattern separation by the GC population and cerebellar learning, but does not influence population sparseness. On a behavioral level, even though locomotor coordination was largely intact in GluA4-KO mice, they were unable to form associative memories during eyeblink conditioning. Together, our results demonstrate that GluA4 is required for reliable synaptic excitation at the cerebellum input layer and for normal cerebellum-dependent associative learning.

### GluA4 Mediates Synaptic Excitation at the Cerebellar Input Layer

Previous studies on GluA4 were mainly performed during development or in young animals (Fuchs et al., 2007; Huupponen et al., 2016; Yang et al., 2011; Zhu et al., 2000; but see: Seol and Kuner, 2015). GluA4 expression decreases during development (Zhao et al., 2019; Zhu et al., 2000). The cerebellar cortex, however, retains high levels of GluA4 in the adult (Keinänen et al., 1990; Monyer et al., 1991; Schwenk et al., 2014), primarily due to the numerous GCs expressing this AMPAR subunit. Here, we show that GluA4-containing AMPARs mediate the majority of excitatory input onto GCs and account for the fast kinetics of MF→GC EPSCs. These results provide additional evidence that the rapid kinetics of GluA4 are crucial for precise timing of postsynaptic spikes (Seol and Kuner, 2015; Yang et al., 2011). Overall, synapses with high GluA4 expression are capable of high-frequency transmission and exhibit strong short-term depression. GluA4 may thus be an integral part of the molecular framework enabling high-frequency information transfer in the mammalian brain.

### Compensations in GluA4-KO Mice

The strongly diminished MF→GC EPSCs in GluA4-KO mice indicate that other AMPAR subunits cannot compensate for the loss of GluA4 (Yan et al., 2013; Yang et al., 2011). We did, however, observe a reduction of tonic inhibition in GluA4-KO GCs, which may enhance information flow through the cerebellar cortex (Hamann et al., 2002). Furthermore, increased NMDAR-EPSCs facilitated spiking of GluA4-KO GCs during high-frequency synaptic input. The larger NMDAR-EPSCs in GluA4-KO are most likely caused by increased glutamate release (Delvendahl et al., 2019), although we cannot exclude postsynaptic contributions. Together, attenuated inhibition and enhanced NMDAR-EPSCs contribute to synaptic information transfer by increasing the excitability of GluA4-KO GCs. However, these adaptations are insufficient for maintaining the fidelity of GC output.

### A Low Number of Active GCs Can Sustain Normal Locomotion

Despite the large reduction of MF→GC transmission, locomotor coordination was not impaired. This may point toward other pathways contributing to sensorimotor coordination or could be due to developmental compensation. It is intriguing to speculate that attenuated inhibition and enhanced non-AMPAR transmission sustain sufficient levels of MF→GC transmission in GluA4-KO animals to spare locomotor activity. The raised GC threshold might also be overcome by activation of multiple MF inputs, which can increase GC spike rates *in vivo* (Ishikawa et al., 2015). MF activity during locomotion is generally dense (Ishikawa et al., 2015; Knogler et al., 2017; Powell et al., 2015), which may activate a sufficient number of GCs for locomotor coordination in GluA4-KO mice. By contrast, the MF representation of the CS used for eyeblink conditioning is most likely not widespread and dense enough to activate a critical fraction of GCs in GluA4-KO mice (Giovannucci et al., 2017; Shimuta et al., 2020). Our finding that in the KO model, GC activity patterns promote faster learning than MF inputs only for high levels of MF activity might thus explain the discrepancy between spared locomotion and absence of associative learning in GluA4-KO mice. These results are in line with previous studies, where impairments of MF→GC or PF→PC synapses did not cause locomotion defects but affected motor learning (Andreescu et al., 2011; Galliano et al., 2013; Peter et al., 2020; Seja et al., 2012). Indeed, a small number of active GCs may be sufficient for basic motor performance (Galliano et al., 2013; Schweighofer et al., 2001).

### The Role of MF→GC Synapses in Associative Learning

Several previous studies investigated PF→PC synaptic function and plasticity in cerebellum-dependent motor learning (Galliano et al., 2013; Grasselli et al., 2020; Gutierrez-Castellanos et al., 2017; Peter et al., 2020; Wada et al., 2007). Afferent sensory information to the cerebellar cortex is first processed at the upstream MF→GC synapse (Billings et al., 2014; Cayco-Gajic et al., 2017), which may contribute to cerebellar learning. Indeed, disturbed AMPAR trafficking at MF→GC as well as other cerebellar synapses in ataxic *stargazer* mice compromises eyeblink conditioning (Hashimoto et al., 1999; Jackson and Nicoll, 2011). Our findings in GluA4-KO mice further support an essential role of MF→GC synapses in cerebellar associative learning. GluA4 is also expressed in Bergmann glia, but deletion of AMPARs in these cells does not affect eyeblink conditioning (Saab et al., 2012). Although we cannot rule out a contribution from other GluA4-containing synapses elsewhere in the brain, associative memory formation during eyeblink conditioning takes place within the cerebellum (Carey, 2011; De Zeeuw and Yeo, 2005; Freeman and Steinmetz, 2011; Heiney et al., 2014; Mauk, 1997; McCormick and Thompson, 1984), where GluA4 is predominantly expressed in GCs. Further, we show here that this AMPAR subunit is crucial for synaptic excitation of cerebellar GCs, but not of Purkinje and Golgi cells. Thus, we conclude that GluA4-containing AMPARs at MF→GC synapses promote cerebellar associative learning. At this synaptic connection, the properties of the GluA4 AMPAR subunit are likely to be important for transmission and processing of the CS, which is conveyed by MF inputs (Albergaria et al., 2018; De Zeeuw and Yeo, 2005; Mauk, 1997).

### GluA4 Supports Computations Underlying Cerebellar Learning

The remaining level of GC excitation in the absence of GluA4 seems sufficient for basic motor performance, but does not allow for associative learning. This indicates that associative memory formation may require more complex levels of GC activity and/or synaptic integration (Albergaria et al., 2018; Carey, 2011; Raymond and Medina, 2018). It is interesting to note that deletion of GluA4 not only caused a strong overall reduction in GC firing frequency, but also severely prolonged the first spike delay upon MF stimulation. GCs are subject to feedback- and feedforward inhibition by GoCs, which narrows the time window for GC integration to ∼5–10 ms (D’Angelo and De Zeeuw, 2009). The longer spike delay in GluA4-KO mice, together with the reduced MF→GC synaptic strength, raises the threshold of GC activation, leading to a loss of information (Billings et al., 2014; Cayco-Gajic et al., 2017). Our findings demonstrate that the higher GC threshold impaired expansion recoding of MF inputs onto the large population of GCs (Albus, 1971; Marr, 1969) (Figure 6G) by effectively reducing the coding space. By contrast, population sparseness of GCs was largely unaffected. Interestingly, the altered GC population coding was sufficient to dramatically impair pattern separation by the cerebellar input layer and learning in downstream circuits. These findings provide additional support for the hypothesis that pattern separation at the cerebellar input layer promotes associative memory formation.

The results we present here provide further evidence for the concept of an AMPAR code (Diering and Huganir, 2018), in which different AMPAR subunits serve different roles for synaptic function and learning. Future studies exploring the possible involvement of GluA4 in synaptic plasticity and other forms of learning will be an important component of our efforts to understand the rules governing cell-type specific expression and function of AMPAR subunits in the CNS.

## Materials and Methods

### Animals

Animals were treated in accordance with national and institutional guidelines. All experiments were approved by the Cantonal Veterinary Office of Zurich (authorization no. ZH206/2016 and ZH009/2020) or by the Portuguese Direcção Geral de Veterinária (Ref. No. 0421/000/000/2015). GluA4-KO mice were kindly provided by H. Monyer (Fuchs et al., 2007). GluA4-KO, GluA4-HET, and wild-type (WT) littermates were bred from heterozygous crosses. Genotyping of GluA4-KO mice was performed as described (Delvendahl et al., 2019). Experiments were performed in male and female mice, 1–5-month-old for slice recordings and 2–5-month-old for behavioral experiments. The animals were housed in groups of 3–5 in standard cages with food and water ad libitum; they were kept on a 12h-light/12h-dark cycle that was reversed for behavioral experiments.

### Electrophysiology

Mice were sacrificed by rapid decapitation either without prior anesthesia or after isoflurane anesthesia in later experiments according to national guidelines. The cerebellar vermis was removed quickly and mounted in a chamber filled with cooled extracellular solution. 300-µm thick parasagittal slices were cut using a Leica VT1200S vibratome (Leica Microsystems, Germany), transferred to an incubation chamber at 35 °C for 30 minutes and then stored at room temperature until experiments. The extracellular solution (artificial cerebrospinal fluid, ACSF) for slice cutting and storage contained (in mM): 125 NaCl, 25 NaHCO_3_, 20 D-glucose, 2.5 KCl, 2 CaCl_2_, 1.25 NaH_2_PO_4_, 1 MgCl_2_, bubbled with 95% O_2_ and 5% CO_2_. For recordings from Golgi cells, the slicing solution consisted of (in mM): 230 sucrose, 25 D-glucose, 24 NaHCO_3_, 4 MgCl_2_, 2.5 KCl, 1.25 NaH_2_PO_4_, 0.5 CaCl_2_, 0.02 D-APV.

Cerebellar slices were visualized using an upright microscope with a 60×, 1 NA water-immersion objective, infrared optics, and differential interference contrast (Scientifica, UK). The recording chamber was continuously perfused with ACSF supplemented with 10 µM D-APV, 10 µM bicuculline and 1 µM strychnine unless otherwise stated. Voltage- and current-clamp recordings were done using a HEKA EPC10 amplifier (HEKA Elektronik GmbH, Germany). Data were filtered at 10 kHz and digitized with 100–200 kHz; recordings of spontaneous postsynaptic currents were filtered at 2.7 kHz and digitized with 50 kHz. Experiments were performed at room temperature (21– 25 °C). Patch pipettes were pulled to open-tip resistances of 3– 8 MΩ (when filled with intracellular solution) from 1.5 mm/0.86 mm (OD/ID) borosilicate glass (Science Products, Germany) using a DMZ puller (Zeitz Instruments, Germany).

The intracellular solution for EPSC and current-clamp recordings contained (in mM): 150 K-D-gluconate, 10 NaCl, 10 HEPES, 3 MgATP, 0.3 NaGTP, 0.05 ethyleneglycol-bis(2-aminoethylether)-N,N,N’,N’-tetraacetic acid (EGTA), pH adjusted to 7.3 using KOH. Voltages were corrected for a liquid junction potential of +13 mV.

We recorded in lobules III–VI of the cerebellar vermis. GC recordings were performed as described previously (Delvendahl et al., 2015, 2019). GC excitability was assessed in current-clamp mode by step current injections of 5–40 pA (duration, 200 ms) from the resting membrane potential. Action potential firing was quantified over the full duration of current injection. GoCs were identified based upon their position in the GC layer, their large capacitance (34.8 ± 3.9 pF, n = 29) and low-frequency of action potential firing (Dieudonne, 1995; Kanichay and Silver, 2008). Spontaneous EPSCs were recorded at a holding potential of – 80 mV. MF→GoC EPSCs were recorded upon stimulation of the white matter or lower GC layer and identified based on the short-latency and rapid kinetics of EPSCs and the absence of short-term facilitation (Kanichay and Silver, 2008) (Figure S1C). PF→GoC EPSCs were recorded upon stimulation of the molecular layer and displayed a longer, variable delay, slower kinetics, and pronounced short-term facilitation (Kanichay and Silver, 2008) (Figure S1D). Recordings from PCs were made similarly; PF→PC EPSCs were recorded upon stimulation of the molecular layer with increasing stimulation voltages (3–18 V) and spontaneous EPSCs were recorded at a holding potential of –80 mV. NMDAR-mediated MF→GC EPSCs were recorded at a holding potential of –80 mV using a Mg-free extracellular solution supplemented with 10 µM NBQX.

To record sIPSCs and tonic inhibitory currents in GCs, we used a CsCl-based intracellular solution containing (in mM): 135 CsCl, 20 TEA-Cl, 10 HEPES, 5 Na_2_phosphocreatine, 4 MgATP, 0.3 NaGTP, 0.2 EGTA, pH adjusted to 7.3 using CsOH (liquid junction potential ∼0 mV). Due to the high intracellular [Cl^−^], IPSCs were recorded as inward currents at a holding potential of –80 mV. 20 µM GYKI-53655 were added to the extracellular solution to block AMPARs. To quantify tonic GABA_A_R-mediated conductance, recordings were made in the presence of 20 µM GYKI-53655 and 1 µM TTX. Following a stable baseline period, 20 µM bicuculline were bath-applied. Tonic conductance was calculated from the difference in holding current before and after bicuculline application, and normalized to cell capacitance to account for account for cell-to-cell variability. GC whole-cell capacitance was not different between genotypes.

To study GC action potential firing upon high-frequency MF stimulation, membrane voltage was maintained at −70 mV (Rothman et al., 2009) by current injection. Similar results were obtained at a membrane potential of −80 mV (Figure S3A). Ten MF stimuli were applied at frequencies of 100–300 Hz. GC spikes were detected using a threshold of −20 mV and the spike frequency was calculated over the entire train duration.

### Western Blotting

Cerebellar tissue was rapidly dissected and frozen at −80°C. Samples were lysed and sonicated in RIPA buffer with protein inhibitors (Roche). Protein concentrations were estimated by BCA assay (Thermo Fisher Scientific) according to manufacturer’s instructions. Protein extracts were then denatured with 2× Laemmi buffer and boiled at 95 °C for 5 min. Samples with equal amount of protein were run on 10% SDS-polyarylamide gel, and then transferred onto PVDF membrane (Thermo Fisher Scientific). Membranes were cut horizontally at appropriate molecular weights and blocked in 5% milk (w/v) in 1× PBS for 1 h on a shaker. Blots were then incubated with primary antibodies: anti-GluA4 (1:100, Santa Cruz Biotechnology) and anti-*β*-Tubulin (1:1,000, DSHB) overnight at 4 °C. To detect transferred proteins, we used horseradish peroxide-conjugated goat anti-mouse antibody (1:1,000, Jackson Immuno Research). Blots were developed with a chemiluminescence detection kit (Thermo Fisher Scientific) and images were acquired with Fusion SL (Vilber). Band intensities were quantified using ImageJ software and normalized to *β*-Tubulin signal.

### MF→GC Modeling

A simple GC adaptive exponential integrate-and-fire model (Brette and Gerstner, 2005) was implemented and run using Neuromatic (Rothman and Silver, 2018) in Igor Pro (Wavemetrics) with a fixed time step of 50 µs. Model parameters were constrained using our experimental data or taken from the literature. To model MF→GC synapses, population averages of 100-Hz AMPAR- and NMDAR-trains were converted into synaptic conductance with direct and spillover components (Rothman et al., 2009). We optimized the model G_AMPA_ and G_NMDA_ parameters to fit the synaptic conductance during 100-Hz trains. Short-term plasticity was implemented using an R*P model (Rothman and Silver, 2018) and parameters were fine-tuned using 100-Hz and 300-Hz population average data for each genotype. Four independent Poisson spike trains were applied to the GC model, with a sum of 10–1000 Hz. Qualitatively similar results were obtained with lower fractions of active MFs. Duration of the spike trains was 1 s and each MF input had a refractory time of 0.6 ms (Ritzau-Jost et al., 2014). Simulations were performed for 37 °C and resting membrane potential was set at −80 mV. The average model spiking frequency was calculated from simulating ten independent runs for each frequency. Plots of GC firing versus MF stimulation frequency were fit with a Hill equation:

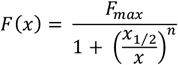

where F_max_ is the maximum firing rate, x_1/2_ the MF stimulation frequency at which F reaches half maximum and n the exponent factor. The offset was calculated taken from x_1/2_ of the fits; the gain was calculated from the slope of the fits between 2% and 60% of the maximum (Rothman et al., 2009).

### Feedforward GC Layer Network Modeling

We adapted the model described in (Cayco-Gajic et al., 2017) to study the effect of GluA4-KO on the function of the cerebellar input layer. In the model, 187 MFs are connected to 487 GCs in a sphere of 80 µm diameter; neuron numbers and connectivity are based on anatomical data. The feedforward model was presented with *n* = 640 different MF input patterns (50 Hz firing, sampled from a Poisson distribution) and GC activity (i.e. spike counts in a 30-ms time window) was extracted. A perceptron algorithm was subsequently used to classify MF and GC activity patterns into 10 random classes (Cayco-Gajic et al., 2017). We compared the published model parameters (Cayco-Gajic et al., 2017) against a model with scaled G_AMPA_ and G_NMDA_ corresponding to the observed relative changes in GluA4-KO GCs (scaling factors 0.2 and 1.2, respectively, Figures 1C and 5B). In addition, the input conductance of the GluA4-KO model was reduced by 0.16 nS (according to Figure 2G). Learning speed, population sparseness and total variance were calculated and analyzed as described in (Cayco-Gajic et al., 2017). In brief, learning speed was analyzed as inverse of the number of training epochs needed to reach a root-mean-square error of 0.2. Population sparseness was calculated as:

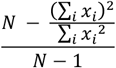

where *N* is the number of GCs and *x*_*i*_ is the ith GC’s spike count. Sparseness was averaged over all *n* activity patterns. Total variance was calculated as the sum of all variances:

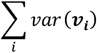

where ***v***_***i***_ is a vector of length *n* consisting of the ith GC’s spike counts for all *n* activity patterns.

### Delay Eyeblink Conditioning

Animals were anesthetized with isoflurane (4% induction and 0.5–1% for maintenance), placed in a stereotaxic frame (David Kopf Instruments, Tujunga, CA) and a head plate was glued to the skull with dental cement (Super Bond – C&B). After the surgery, mice were monitored and allowed at least 2 days of recovery.

All eyeblink conditioning experiments were run on a motorized treadmill, as described in previous work (Albergaria et al., 2018). Briefly, head-fixed mice were habituated to the motorized behavioral setup for at least 2 days prior to training. The speed of the treadmill was set to 0.12 m/s using a DC motor with an encoder (Maxon). After habituation, each training session consisted of 100 trials, separated by a randomized inter trial interval of 10–15 s. In each trial, CS and US onsets were separated by a fixed interval of 300 ms and both stimuli co-terminated.

For all training experiments, the unconditioned stimulus (US) was an air-puff (30–50 psi, 50 ms) controlled by a Picospritzer (Parker) and was adjusted for each session of each mouse so that the unconditioned stimulus elicited a full eye blink. The CS had a 350 ms duration and was either i) a white light LED positioned ∼3 cm directly in front of the mouse or ii) a piezoelectric device placed ∼0.5 cm away from the left vibrissal pad.

Eyelid movements of the right eye were recorded using a high-speed monochromatic camera (Genie HM640, Dalsa) to monitor a 172 × 160 pixel region, at 900 fps. Custom-written software using LabVIEW, together with a NI PCIE-8235 frame grabber and a NI-DAQmx board (National Instruments), was used to trigger and control all the hardware in a synchronized manner.

### Locomotion Analysis

We used the LocoMouse system to quantify overground locomotion (Machado et al., 2015). Animals run back and forward between two dark boxes, in a glass corridor (66.5 cm long and 4.5 cm wide with a mirror placed at 45° under). Individual trials consisted of single crossings of the corridor. A single high-speed camera (AVT Bonito, 1440 × 250 pixels, 400 fps) recorded both bottom and side views of walking mice during the trial. Animals were acclimated to the overground setup for several sessions before data collection; no food or water restriction or reward was used.

Homozygous GluA4-KOs ranged in size from 16–37 g. Size-matched littermate HET (18–38 g) and WT animals (16–38 g) were selected as controls (Machado et al., 2015). Ten to twenty-five corridor trials were collected in each session for five consecutive days. An average of 1207 ± 203 strides were collected per homozygous GluA4 mouse (310 ± 51 strides per animal per paw), 856 ± 342 strides per heterozygous GluA4 mouse (219 ± 94 strides per animal per paw) and 840 ± 278 strides per WT mouse (218 ± 70 strides per animal per paw) were collected.

## Data Analysis

Electrophysiological data were analyzed using custom-written routines or Neuromatic (Rothman and Silver, 2018) in Igor Pro (Wavemetrics, USA). EPSC amplitudes were quantified as difference between peak and baseline. Decay kinetics were analyzed by fitting a bi-exponential function to the EPSC decay time course. To detect spontaneous or miniature postsynaptic currents, we used a template-matching routine (Rothman and Silver, 2018). Detected events were analyzed as described above. Only cells with > 10 detected spontaneous events were included for amplitude quantification.

Simulation results were analyzed using Igor Pro and Python. To calculate the van Rossum distance, pre- and postsynaptic spike trains were convolved using a 10-ms exponential kernel. The area of the squared difference was then multiplied by the inverse of the kernel time constant to give the van Rossum distance. Spike synchronization was calculated using PySpike (Mulansky and Kreuz, 2016).

For eyeblink conditioning, video from each trial were analyzed offline with custom-written software using MATLAB (MathWorks). The distance between eyelids was calculated frame by frame by thresholding the grayscale image of the eye and extracting the count of pixels that constitute the minor axis of the elliptical shape that delineates the eye. Eyelid traces were normalized for each session, ranging from 1 (full blink) to 0 (eye fully open). Trials were classified as CRs if the eyelid closure reached at least 0.1 (in normalized pixel values) and occurred between 100 ms after the time of CS onset and the onset of US.

Analysis of locomotion data was performed in Matlab 2012b and 2015a. Paw, nose and tail tracks (x,y,z) were obtained from the LocoMouse tracker (Machado et al., 2015) (https://github.com/careylab/LocoMouse). All tracks were divided in stride cycles and the data was sorted into speed bins (0.05 m/s bin width). Gait parameters (individual limb movements and interlimb coordination) were calculated as follows:

### Individual limb

<u>Walking speed</u>: x displacement of the body center during that stride divided by the stride duration

<u>Stride length</u>: x displacement from touchdown to touchdown of single limb

<u>Stride duration</u>: time between two consecutive stance onsets <u>Cadence</u>: inverse of stride duration

<u>Swing velocity</u>: x displacement of single limb during swing phase divided by swing duration

<u>Stance duration</u>: time in milliseconds that foot is on the ground during stride

<u>Duty factor</u>: stance duration divided by stride duration <u>Trajectories</u>: (x,y,z) trajectories were aligned to swing onset and resampled to 100 equidistant points using linear interpolation. Interpolated trajectories were then binned by speed and the average trajectory was computed for each individual animal and smoothed with a Savitzky-Golay first-order filter with a 3-point window size.

<u>Instantaneous swing velocity</u>: the derivative of swing trajectory

### Interlimb and whole-body coordination

<u>Stance phase</u>: relative timing of limb touchdowns to stride cycle of reference paw (FR). Calculated as: (stance time − stance time _reference paw_) / stride duration.

<u>Base of support</u>: width between the two front and two hind paws during stance phase

<u>Body y displacement</u>: y displacement of the body center during that stride

<u>Supports</u>: Support types were categorized by how many and which paws were on the ground, expressed as a percentage of the total stride duration for each stride. Paw support categories include 3-paw, 2-paw diagonal, 2-paw other/non-diagonal (homolateral and homologous), and 2-paw front (only) supports.

<u>Double support for each limb</u> is defined as the percentage of the stride cycle between the touch down of a reference paw to lift-off of the contralateral paw. Because at higher speeds (running), the opposing limb lifts off before the reference paw touches down, we included negative double support by looking backwards in time, up to 25% of the stride cycle duration. Positive values of double support indicate that contralateral lift-off occurred after reference paw touch down, and negative values indicate that contralateral lift-off occurred before reference paw touch down. Note that front paw double support percentages include 2-paw front (only) support patterns as well as 3- and 4-paw support patterns in which both front paws were on the ground.

<u>Tail and nose phases</u>: For each speed bin we correlate the stridewise tail and nose trajectories with the trajectory given by the difference between the forward position of the right paw and the forward position of the left paw (also normalized to the stride). The phase is then calculated by the delay in which this correlation is maximized.

<u>Tail and nose peak-to-peak amplitude</u>: the change between peak (highest amplitude value) and trough (lowest amplitude value) in y or z during a stride duration.

<u>Variability</u>: All variability analyses were based on coefficients of variation (CV).

### Principal Component and Linear Discriminant Analyses

The locomotor dataset consisted of a matrix of 45 features for each mouse and speed bin. Many gait features are highly correlated with speed (Machado et al., 2015), so to avoid inter-variable correlation and overfitting we first performed PCA. The first 10 principal components explained more than 85% of the variance and the data projected onto these 10 principal components were used as input to the LDA (Machado et al., 2020). LDA output is displayed separately for each speed bin to verify that the pattern of differences across groups was captured across all walking speeds.

### Statistical Analysis

Electrophysiological data were analyzed using ANOVA and two-tailed t-tests. Statistical testing was performed in R. Data in figures are presented as mean ± standard error of the mean (SEM). Eyeblink conditioning data were analyzed using t-tests. Locomotion data were analyzed in Matlab with the Statistics toolbox. An independent samples t-test was used to test for differences in walking speed distributions (Figure S6A). For all other gait parameters, analysis was performed on animal averages binned by speed using mixed effects models (Bates et al., 2015). Fixed-effects terms included speed and genotype; animals were included as random terms. Table S1 reports effects of genotype for all locomotor measurements as F statistics from mixed ANOVAs with Satterthwaite degrees of freedom correction. No corrections were made for multiple comparisons.

